# TCR-H: Machine Learning Prediction of T-cell Receptor Epitope Binding on Unseen Datasets

**DOI:** 10.1101/2023.11.28.569077

**Authors:** Rajitha Rajeshwar T., Omar Demerdash, Jeremy C. Smith

**Affiliations:** UT/ORNL Center for Molecular Biophysics, Oak Ridge National Laboratory, Oak Ridge, Tennessee 37831-6309, United States; Department of Biochemistry and Cellular and Molecular Biology, University of Tennessee, Knoxville, Tennessee 37996-1939, United States; Biosciences Division, Oak Ridge National Laboratory, Oak Ridge, Tennessee 37831-6309, United States

## Abstract

AI/ML approaches to predicting T-cell receptor (TCR) epitope specificity achieve high performance metrics on test datasets which include sequences that are also part of the training set but fail to generalize to test sets consisting of epitopes and TCRs that are absent from the training set, i.e., unseen. We present TCR-H, a supervised classification Support Vector Machines model using physicochemical features trained on the largest dataset available to date using only experimentally validated non-binders as negative datapoints. TCR-H exhibits an area under the curve of the receiver-operator characteristic (AUC of ROC) of 0.87 for epitope ‘hard splitting’ (i.e., on test sets with all epitopes unseen), 0.92 for TCR hard splitting and 0.89 for ‘strict splitting’ in which neither the epitopes nor the TCRs in the test set are seen in the training data. TCR-H may thus represent a significant step towards general applicability of epitope:TCR specificity prediction.

## Introduction

Cytotoxic T-cells are central to the adaptive immune response. Critical to adaptive immune system activation is the specific binding of T-cell receptors (TCR) to peptide epitopes presented by the Major Histocompatibility Complex (MHC) on the surface of antigen-presenting cells. The remarkable diversity of TCRs, estimated at ∼10^15^-10^61^ (Dash et al., 2017), results from a combinatorial explosion of genetic recombination possibilities of somatic-cell DNA encoding V (variable), D (diversity), and J (joining) segments(Bradley & Thomas, 2019; Rudolph et al., 2006). T-cells are thus capable of recognizing a large variety of epitopes, that can be exogenous or derived from endogenous mutated proteins (Tippalagama et al., 2023).

A significant goal in immunology is the reliable prediction of which epitopes bind to which T-cell receptors. Knowing this would aid in the design and development of vaccines and immunotherapies and would help us understand how the immune system distinguishes between self and non-self antigens. Furthermore, knowledge of TCR:epitope specificity can be used in disease diagnosis, prognosis, and monitoring disease progression, which is particularly important in the context of infectious diseases, where tracking T-cell responses can provide insights into the immune system’s ability to recognize and control pathogens.

The vast diversity in TCR sequences and potential epitopes makes it challenging to develop a generalized computational model for TCR:epitope binding prediction. Moreover, prediction is further complicated by the fact that TCRs can exhibit cross-reactivity, recognizing multiple epitopes. One approach, in principle, to solving the TCR:epitope specificity problem would be to develop accurate methods for predicting the 3D structures of TCR:epitope:MHC ternary complexes and then to use the results to predict binding strength based on the physics involved. However, predicting these 3D structures and interactions accurately is challenging (Bradley 2023). The 3D interaction between TCRs and epitopes bound to MHC is highly variable and small changes in epitope sequence can significantly impact TCR recognition. Conformational flexibility of TCRs and the epitopes they recognize further complicates this approach.

In recent work we demonstrated how 3D structure can be incorporated into computational methods to improve prediction of peptide binding to MHC (Aranha et al., 2020), and how 3D MHC pocket similarity determines peptide binding strength (Yue et al., 2023). Furthermore, we and others have shown that certain 3D interactions are commonly found in epitope binding to TCRs(Rajeshwar & Smith, 2022). We therefore postulated that TCR:epitope recognition algorithms would benefit from further consideration of physicochemical properties determining recognition.

TCRs are heterodimers consisting of two chains: α and β. Binding primarily occurs through three complementarity-determining regions (CDRs) found on each of these chains. Although in 3D structures of TCR:epitope:MHC ternary complexes significant contacts are made outside the CDR3 region, this region most directly interacts with the peptide epitope, with CDR3β, in particular, forming direct contacts with the epitopes and MHC and displaying particularly high sequence diversity (Dai et al., 2008; Garcia et al., 2009; Morris & Allen, 2012; Szeto et al., 2021, Rajeshwar & Smith, 2022)). Hence, in computational approaches CDR3β has commonly been taken as an approximation to the specificity-determining TCR sequence.

With the increasing availability in publicly available data resources of CDR3β and epitope sequencing data from high-throughput techniques (Jokinen et al., 2021), AI and ML approaches have been able to be used to predict TCR CDR3β binding to epitopes presented by MHC class 1 (MHC-I) (Hudson et al., 2023). These tools apply methods ranging from relatively simple ML models such as Random Forest and clustering (Chronister et al., 2021; Dash et al., 2017; Jokinen et al., 2021; Pham et al., 2023) to various forms of deep learning-based AI techniques, including convolutional and recurrent neural networks (Bravi et al., 2023; Cai et al., 2022; Darmawan et al., 2023; Gao, Gao, Li, et al., 2023; Jiang et al., 2023; Jokinen et al., 2023; Myronov et al., 2023; D. Wang et al., 2023; Z. Wang & Shen, 2023; Weber et al., 2021; Zhang et al., 2023).

Many of the above AI/ML methods achieve impressive results on ‘random split’ test sets that include TCR and epitope sequences that are also included in the training set. However, for general applicability a model must be able to predict binding for a “strict split” with unseen TCRs and epitopes, i.e., in which neither the TCRs nor the epitopes tested are in the training set. Success in this endeavor would perhaps indicate an algorithm has been discovered that has started to learn the general principles of TCR:epitope recognition and would be a major step towards a “holy grail” of immunology - the accurate computational mapping of all TCRs to their cognate epitopes.

The strict split goal can be approached stepwise. In a first step, datasets can be derived with an epitope “hard split”, i.e., in which the test set contains only unseen epitopes, and a TCR hard split in which the test set contains only unseen CDR3βs. When using an epitope hard split, current methods mostly fail, with scores falling to almost random i.e., with the area under the curve of the receiver-operator characteristic (AUC) ∼0.5. (Dens et al., 2023; Grazioli et al., 2022). TCR hard splitting has been performed in (Cai et al., 2022) with AUC values of 0.77 or below. Strict splitting attempts have also been published with AUC values ranging between 0.5 and 0.7 (Korpela et al., 2023; Weber et al., 2021)

Given the above considerations, we attempted to address the unseen epitopes and TCRs problem by developing a supervised binary classification ML model incorporating sequence-based physicochemical descriptors of CDR3β and epitopes and using the largest dataset of binding and non-binding data available to date. Our model possesses two noteworthy characteristics: the use of only experimentally validated non-binders as negative datapoints in the training and testing sets and the use of a highly diverse set of physicochemical features calculated over entire peptide sequences.

Existing machine and deep learning methods vary in terms of the datasets used, mostly relying on randomly generated data as the negative/non-binding dataset. However, any given randomly generated sequence assumed to not bind might actually bind. Furthermore, it has been shown that the use of randomly generated sequences for negative binders leads to overestimation of model accuracy (Dens et al., 2023; Grazioli et al., 2022). Therefore, we include in our dataset only negatives that have been experimentally validated as non-binders.

The choice of supervised ML over deep learning was motivated in part by the ability of the model to be based on explicit features posited to be important for binding, which in turn may aid future model explainability and rationality. Furthermore, it has also been shown that the use of features that lend themselves to ML models leads to better performance in TCR specificity prediction (Pham et al., 2023). Most ML models have been sequence-based, encoding sequences using either the BLOSUM substitution matrix (Cai et al., 2022; Darmawan et al., 2023; Gao, Gao, Li, et al., 2023; Pham et al., 2023) and/or Atchley factors(Atchley et al., 2005). The BLOSUM matrix applies a score for amino-acid substitutions while Atchley factors are multidimensional, composite features for each amino acid derived using unsupervised ML on primarily physicochemical features (Jiang et al., 2023; Lu et al., 2021; Moris et al., 2021). Here, in contrast we use physicochemical features calculated over entire epitope and CDR3β sequences i.e., not just individual amino-acid residues.

Various ML methods are tested: Random Forest (RF), Gradient Boosting trees (GBT), eXtreme Gradient Boosting (xGBT) and Support Vector Machines (SVM). The best performing, an SVM model, which we name TCR-H, exhibited area under the receiver-operator characteristic curve (AUC of ROC) metrics of 0.87, 0.92 and 0.89 for the epitope hard split, TCR hard split and the epitope/TCR strict split, respectively. These are significant improvements on previously reported values and TCR-H may thus represent a noteworthy step towards general applicability of computational prediction of epitope:TCR specificity in biology and medicine.

## Results and Discussion

Details of the dataset used, which comprises ∼250,000 datapoints, the featurization and the ML models tested are given in Methods. Our binary classification is the prediction of binding or non-binding. The performance of the models was evaluated in terms of the AUC of ROC, the accuracy, the precision (the ratio of true binders to the total number predicted to be binders, also called positive predictive value), the recall (also called the sensitivity or true positive rate), the specificity (true negative rate), and the F1 score (see SI for definitions). The AUC of ROC, which is most commonly reported, illustrates the trade-off between the recall (true positive rate) and the specificity (true negative rate), quantifying the ability of the model to distinguish between the two classes, where a classifier with perfect performance yields an AUC of ROC of 1.0, while a random classifier would yield an AUC of 0.5 or less.

We first present ML performance results on epitope hard split data, i.e., in which the only epitopes in the test set are unseen. The resulting performance metrics on the test set for RF, GBT, xGBT and SVM are presented in Table 1. Taking all the performance metrics into consideration, it is clear that the ensemble models RF, GBT and xGBT do not perform particularly well, especially for AUC of ROC and specificity. Only in terms of recall and F1-score do these models perform somewhat decently; however, it should be noted that the F1, a composite metric that is a function of both precision and recall, is elevated here only by the recall, with precision being relatively poor. In contrast, for the model trained with SVM, performance is much better across the board, with the AUC of ROC, accuracy, and precision—all poor in the ensemble tree models—improving dramatically. The AUC of ROC is 0.80 (Figure S1), an improvement on previously reported efforts.

**Table 1.** Comparison of ML models for epitope hard split test set. TP=number of true positives; TN=number of true negatives; FP=number of false positives; FN=number of false negatives. TCR-HE has correlations removed.

| Model | AUC of ROC | TP | TN | FP | FN | Accuracy | Precision | Recall | Specificity | F1 |
| --- | --- | --- | --- | --- | --- | --- | --- | --- | --- | --- |
| XGB | 0.51 | 31859 | 654 | 21817 | 104 | 0.597 | 0.593 | 0.996 | 0.029 | 0.744 |
| GBT | 0.54 | 31248 | 2329 | 20142 | 715 | 0.617 | 0.608 | 0.977 | 0.104 | 0.749 |
| RF | 0.5 | 31963 | 0 | 22471 | 0 | 0.587 | 0.587 | 1.00 | 0.0 | 0.739 |
| SVM | 0.80 | 30403 | 14593 | 7878 | 1560 | 0.826 | 0.794 | 0.951 | 0.649 | 0.865 |
| TCR-HE | 0.869 | 29567 | 18297 | 4174 | 2396 | 0.879 | 0.876 | 0.925 | 0.814 | 0.9 |

In an attempt to further improve the SVM performance, we calculated correlations between pairs of features. We found that by removing one of any pair that had a correlation > 0.8 (we selected the feature to be removed from the correlated pair at random) improved the model’s performance leading to a model (named “TCR-HE”) with an AUC of ROC of 0.87 (Fig. 1A) as well as improving most other statistical metrics. The model has very high sensitivity, i.e., a large fraction of the true positives are predicted positive. The specificity (true negative rate) is also high, although perhaps not quite as impressive. To further test the robustness of the model’s performance, we conducted three other epitope hard splits while introducing imbalanced numbers of negative and positive data points. The resulting performance metrics are detailed in the supporting information (Table S2). The AUC for these scenarios ranged between 0.72 and 0.853, maintaining high, if not quite as high, performance.

**Figure 1.**
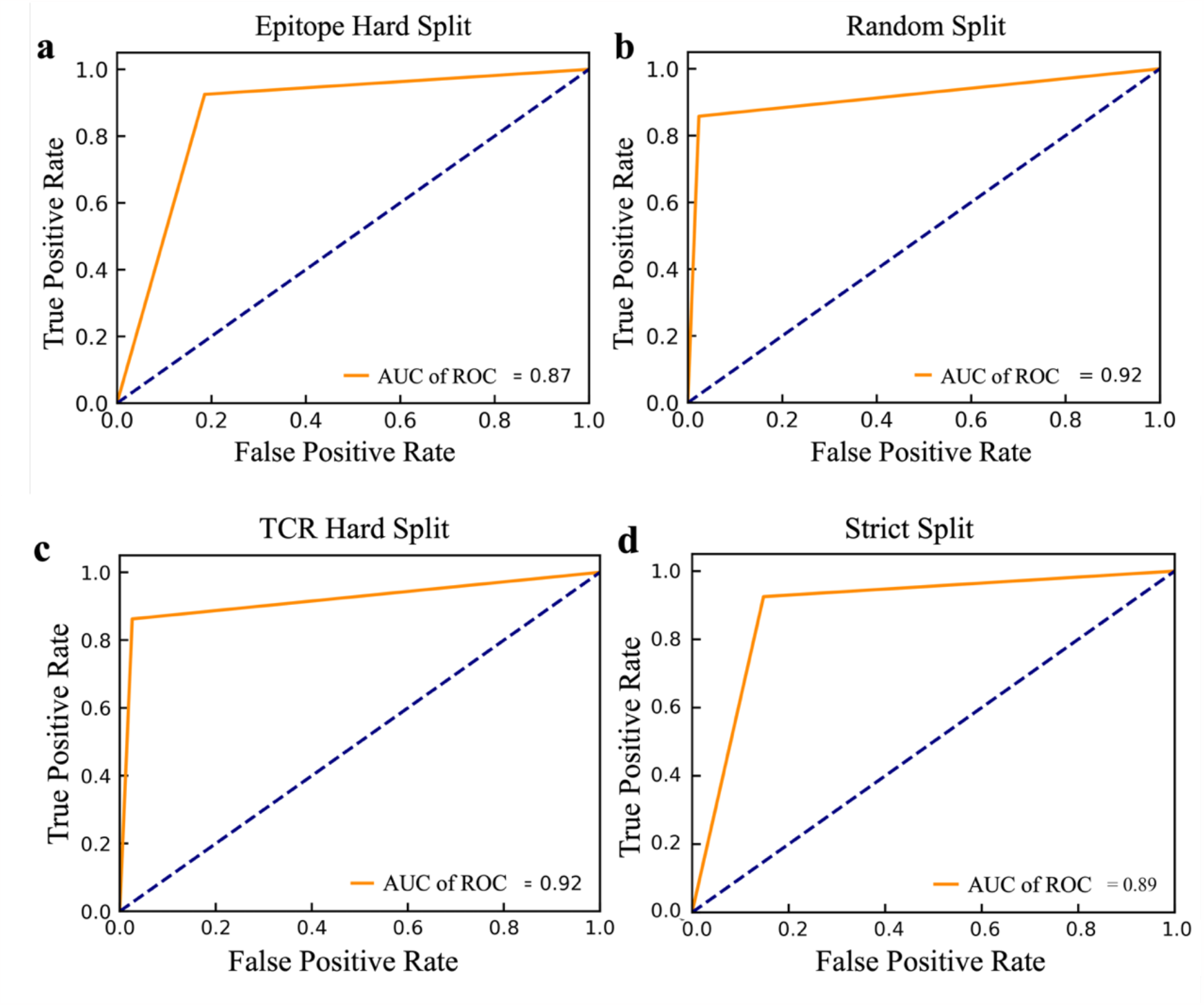
AUC of ROC on the independent test set for the TCR-H model with uncorrelated features trained with a) TCR-HE epitope hard split b) TCR-HRS random split. C) TCR-Hβ hard split of the TCRs. D) TCR-HβE strict split .

Next, we trained the model using SVM and the set of uncorrelated features using a hard split in the TCR CDR3βs (a model referred to as TCR-Hβ). The results on the independent test set, provided in Table 2, show that again the model performs very well, with an AUC of ROC of 0.92 and other metrics also being high. Again, to test dataset robustness we conducted three additional hard splits for TCR CDR3β and also three for the strict split (Table S2). The performance metrics consistently demonstrated an AUC of ROC of 0.92 for the different TCR hard splits. This model appears to be particularly stable to dataset variation, and this may arise from there being ∼150k unique TCR sequences in the dataset as compared to ∼800 unique epitope sequences.

**Table 2.** Performance metrics of TCR-H trained and tested on TCR hard split (TCR-Hβ), strict split (TCR-HβE) and random split (TCR-HRS).

| Model | AUC of ROC | TP | TN | FP | FN | Accuracy | Precision | Recall | Specificity | F1 |
| --- | --- | --- | --- | --- | --- | --- | --- | --- | --- | --- |
| TCR-Hβ | 0.918 | 18760 | 28780 | 766 | 2994 | 0.93 | 0.96 | 0.86 | 0.97 | 0.91 |
| TCR-HβE | 0.888 | 29570 | 19143 | 3328 | 2393 | 0.89 | 0.898 | 0.92 | 0.85 | 0.91 |
| TCR-HRS | 0.917 | 18382 | 28771 | 691 | 3045 | 0.93 | 0.96 | 0.86 | 0.98 | 0.91 |

Finally, we report metrics on the strict split, TCR-HβE, which gives an AUC of ROC of 0.89 with the other metrics again high. The strict splits with imbalanced data yielded AUC of ROC values ranging between 0.71 and 0.83 (Table S2), again with excellent, if not quite as good, performance.

For completeness, and to compare against what has been typically done in prior predictive model development studies, we also performed a benchmark of TCR-H using the commonly employed random split (TCR-HRS). In a random split the entire dataset is split randomly into training and test sets. Because the dataset contains instances where a TCR or epitope can be represented multiple times, a random split of the data can lead to the same epitope or TCR being in both the training and test sets. For TCR-HRS, the AUC of ROC of TCR-H model is 0.92 (Table 2) which is comparable to previously-published methods. Additionally, three other random splits employed also showed an AUC of ROC of 0.92, again indicating model robustness.

### Comparison with previous models

The field of computational epitope-TCR binding prediction is very active and rapidly evolving. However, it is incumbent on us to compare the performance of TCR-H with models hitherto reported in the literature (Cai et al., 2022; Gao, Gao, Fan, et al., 2023a; Gao, Gao, Li, et al., 2023; Jiang et al., 2023; Lu et al., 2021; Montemurro et al., 2021; Moris et al., 2021; Pham et al., 2023). Metrics other than the AUC of ROC are important in potential applications of these methodologies but are not always calculated. However, data exist allowing us to compare the performance of TCR-HE with some reported data for epitope hard-split AUC of ROC, precision, and recall . Fig. 2 shows metrics for various methods taken from Cai et al. (2022) all of which were evaluated on the same independent epitope hard split data set, together with the present TCR-HE results. TCR-HE exhibits higher values of all three metrics. Some methods, including Pan-Peptide and epiTCR, have in some epitope hard-split tests shown significant improvement over random, with AUC of ROC scores of ∼0.75. Notwithstanding, TCR-HE performed significantly better. Furthermore, the pitfalls of negative data bias in the TCR epitope specificity challenge have recently been emphasized, with, for example, the performance of Pan-peptide reduced to random (AUC of ROC = 0.49) depending on the method of choosing the negative data points (Dens et al., 2023).

**Figure 2.**
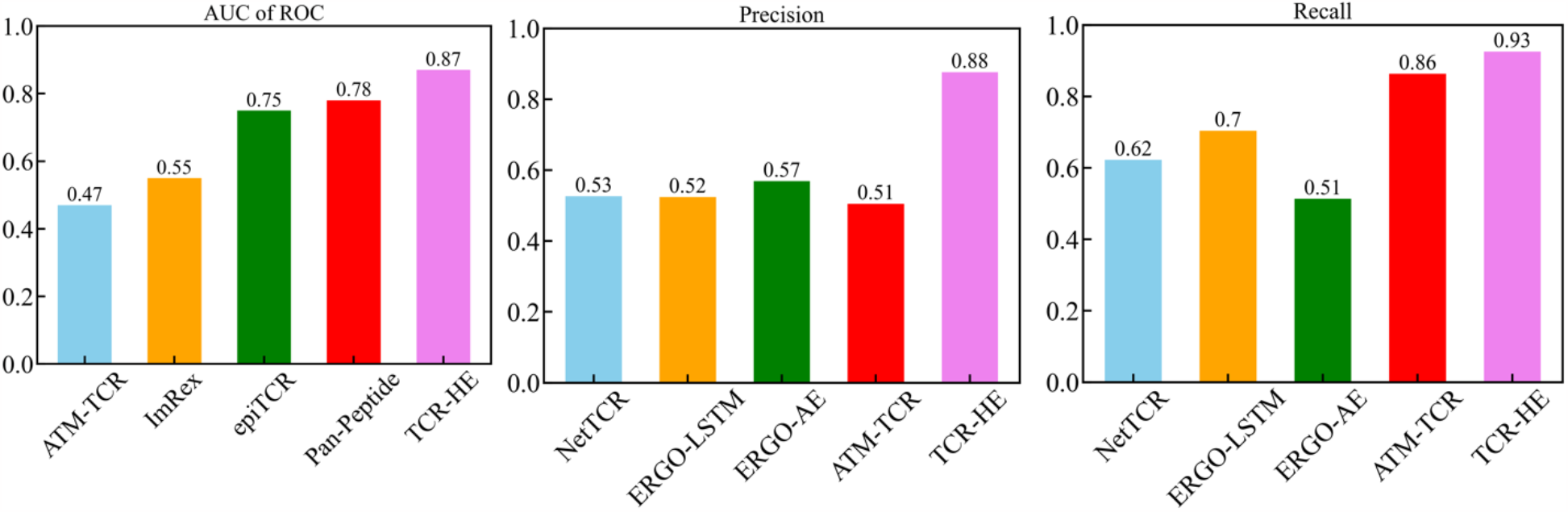
AUC of ROC, Precision and Recall of TCR-HE compared with that of previously reported epitope hard split models (Cai et al. (2022); Moris et al., 2021; Pham et al., 2023 ;Gao, Gao, Fan, et al., 2023).

Published data on TCR hard splitting and strict splitting are rarer. However, Cai et al 2022 report TCR hard split metrics for four models (NetTCR, ERGO-LSTM, ERGO-AE and ATM-TCR), with AUC of ROC varying between 0.72 and 0.77, recall between 0.71 and 0.77 and precision between 0.63 and 0.70, again lower than the values obtained with TCR-Hβ (Table 2) whereas TITAN (Weber et al 2021) achieved an AUC of ROC of 0.87 using a semi-frozen pretrained model with augmentation. A strict split employed by TITAN (Weber et al 2021) achieved an AUC of ROC of 0.62.

## Conclusion

Building on our previous work on 3D structural and physicochemical approaches to studies of MHC:epitope:TCR interactions, we present here an ML model, TCR-HβE, capable of predicting TCR and epitope binding for the ‘strict split’ case in which neither the epitopes nor the TCRs were included in data used to train the model. Our approach to this was stepwise, in which the performance of TCR-H was first tested on hard splits of epitopes (TCR-HE) and TCRs (TCR-Hβ). Both of these produced excellent metric results that were robust to training/testing dataset variation with the TCR-Hβ being particularly impressive. The strict split TCR-HβE achieved an AUC of ROC of 0.89, with other metrics such as precision and recall also excellent.

AI/ML methods can be notoriously fickle and subject to biases in the data sets used for training and testing. We therefore paid special attention to quality of the datasets used. In particular we only considered experimentally validated data, as opposed to using negative data that is randomly generated. Notwithstanding, testing of a given model on different specific epitope hard split data sets can lead to significantly different AUC values. For example, Fig. S3 shows significant dataset dependency of epitope hard split AUC of ROC data with previously-reported models. Here, we made an effort to examine the performance of the TCR-H class of models with dataset variation. We find good performance even when the test data are burdened with significant positive and negative data imbalance. However, we believe that, although the dataset used has >200,000 entries, the approach taken here, as for most models, will need continual further testing and refining. In particular experimental addition to the number of unique epitopes would be expected to further improve model performance. Furthermore, as stated in Montemurro et al., 2021 there is room for improvement in the accuracy of the negative data generated.

The good performance of the TCR-H models suggests that supervised SVM, when applied to a feature set consisting of physicochemical features derived from whole sequences of TCR CDR3βs and epitopes, may be able to capture complex feature dependencies and interdependencies underlying accurate binding classification, It is interesting that ensemble tree-based methods did not perform as well as SVM, suggesting that to achieve accurate binding prediction, a complex function of the full set of features is advantageous, as opposed to an ensemble of simple, nonparametric “functions”, as represented by the simple individual trees.

For general applicability, a TCR:epitope recognition model must be able to predict binding for TCRs and epitopes for any given TCR:epitope pair. This should include cases in which neither the TCRs nor the epitopes queried are in the training set nor are close to members in the training set, i.e., the “strict split” scenario. The success in this endeavor reported here may indicate that algorithms are learning general principles of TCR:epitope recognition. We see TCR-H as a step in this direction, and towards making the vast universe of potential epitope and TCR sequences amenable to computational prediction of functional TCR/epitope pairs.

## Methodology

### Dataset

The dataset used comprises positive binding data curated for human MHC class 1 from the IEDB, VDJdb, and McPAS-TCR databases, and negative binding data from IEDB and 10X Genomics assays, sourced from TChard (Grazioli et al., 2022). The dataset includes CDR3β sequence lengths ranging from 9 to 23 residues, while the epitope sequences have lengths shorter than 16. The total dataset consists of 147,069 negative and 107,376 positive datapoints and was divided into training and test data sets of 80 and 20 percent, respectively.

Two different types of data split were considered 1) Hard Split 2) Random Split. In the epitope hard split the test set consisted of only unseen epitopes. The epitope hard split training data set in Table 2 (total: 200,011) consisted of 124,598 negative datapoints and 75,413 positive data whereas the test dataset (total: 54,434) consisted of 22,471 negative data and 31,963 positive data points, with 65 unseen epitopes. It should be noted that most studies do not report the number of unseen epitopes in the test dataset. For the TCR hard split in Table 2, again the training and test data were split approximately in the ratio of 80:20. In the test dataset with unseen TCR CDR3βs, out of 51,300 data points there were 21,754 positive data and 29,546 negative data. In the strict split, the training data consisted of 65826 positive data and 83697 negative data points whereas test dataset consisted of 22,471 negative data and 31,963 positive data. The training and test data sets are detailed in the supporting information (Table in SI).

### Feature Set

All sequence-based properties of the CDR3β loops and the epitopes were calculated using the Peptides Python package [https://github.com/althonos/peptides.py] (Osorio et al., 2015), resulting in 96 features each for the CDR3β and epitope sequences. A complete list of the features used is given in Table 4. In the feature set used to train the ML models, some features differ substantially in magnitude from one another. In training ML models whose optimization is gradient-based, the presence of features differing by an order of magnitude or more can pose a numerical challenge and yield erroneous results. To avoid this problem, features were normalized, or scaled, so that they were all within the same order of magnitude. To achieve this, the *preprocessing*.*scale* function in Scikit Learn was used, which centers features to the mean and normalizes them to unit variance.

**Table 4:** Sequence-based properties used as features (96 each for epitope and CDR3b) in the present study, the numbers in the parentheses represents the number of features corresponding to given property.

|  |  |
| --- | --- |
| BLOSUM indices | 10 |
| Cruciani properties | 8 |
| FASGAI vectors | 6 |
| Kidera factors | 10 |
| MS-WHIM scores | 3 |
| PCP descriptors | 5 |
| Physical descriptors | 2 |
| ProtFP descriptors | 8 |
| Sneath vectors | 4 |
| svger_descriptors | 10 |
| ST-scales | 8 |
| T-scales | 5 |
| VHSE-scales | 8 |
| Z-scales | 5 |
| Boman | 1 |
| Charge | 1 |
| Hydrophobic Moment | 1 |
| Hydrophobicity | 1 |
| Isoelectric Point | 1 |
| Molecular Weight | 1 |
| m/z(mass/charge) | 1 |

### Machine Learning Methods

We examined four ML methods: Random Forest (RF), Gradient Boosting trees (GBT), eXtreme Gradient Boosting and Support Vector Machines (SVM). All models were trained using the Scikit Learn application programming interface.

Random Forest (RF) is an ensemble learning method, *i*.*e*., in which the final model is a composite of multiple individual models. RF operates by constructing multiple simple decision trees during training. Each tree in the forest is constructed using a random subset of the features. The final prediction is based on the aggregated votes of individual trees. Here, the RF model was built using default hyperparameters and scaled features. Gradient-Boosting Trees (GBT) is another ensemble method we tested. In contrast to RF, whose trees are constructed independently of one another during model training, GBT successively augments the model in each iteration with a tree such that it minimizes a loss function representing the discrepancy between the training targets and the corresponding model predictions. eXtreme Gradient Boosting (XGB) is similar to GBT in that it is based on minimizing a loss function with each tree that is successively added to the ensemble of trees, but instead of optimizing using just the gradient, it makes use of the Hessian as well, in effect optimizing using a Newton-Raphson type of approach.

In classification, SVM aims to find the hyperplane that best separates the data into different classes, maximizing the margin between the classes. SVM can be trained on the original set of features (corresponding to a linear SVM model) or a projection of the original set of features into a non-linear space using different kernel functions. We used the Radial Basis Function (RBF) kernel, also known as the Gaussian kernel, with the default hyperparameters. The prediction model was built with scaled features. Variation of the hyperparameters did not improve model performance.

## Acknowledgements

This research used resources of the Compute and Data Environment for Science (CADES) at the Oak Ridge National Laboratory, which is supported by the Office of Science of the U.S. Department of Energy under Contract No. DE-AC05-00OR22725.

## Supporting Information

### Dataset Details

In Table S1 we present the categorized numbers of data in the various datasets used.

**Table S1:** Categorized dataset numbers.

Epitope Hard Split:
| Training dataset |  |  | Test dataset |  |  |
| --- | --- | --- | --- | --- | --- |
| Positive data | Negative data | Unique peptides | Positive data | Negative data | Unique peptides |
| 75413 | 124598 | 790 | 31963 | 22471 | 65 |

TCR Hard Split:
| Training dataset |  |  | Test dataset |  |  |
| --- | --- | --- | --- | --- | --- |
| Positive data | Negative data | Unique CDR3βs | Positive data | Negative data | Unique CDR3βs |
| 85621 | 117523 | 124098 | 21754 | 29546 | 31025 |

Strict Split:
| Training dataset |  |  |  | Test dataset |  |  |  |
| --- | --- | --- | --- | --- | --- | --- | --- |
| Positive data | Negative data | Unique CDR3βs | Unique peptides | Positive data | Negative data | Unique CDR3βs | Unique peptides |
| 65826 | 83697 | 105011 | 739 | 31963 | 22471 | 50112 | 65 |

### Performance Metric Definitions

The performance model metrics are defined as follows:

Accuracy (ACC) is the proportion of correctly classified instances:

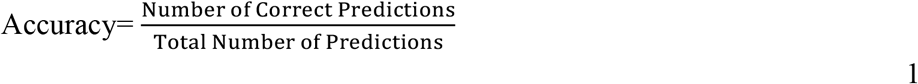

Precision (also called Positive Predictive Value) is the proportion of true positives among instances predicted as positive:

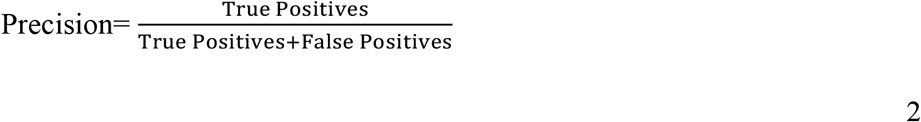

Recall (also called Sensitivity or True Positive Rate) is the proportion of true positives among actual positive instances:

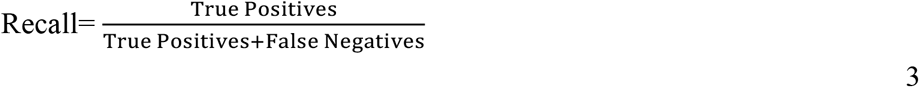

F1 Score is the harmonic mean of precision and recall:

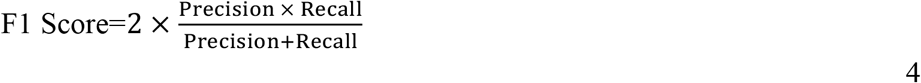

Specificity (True Negative Rate) is the proportion of true negatives among actual negative instances:

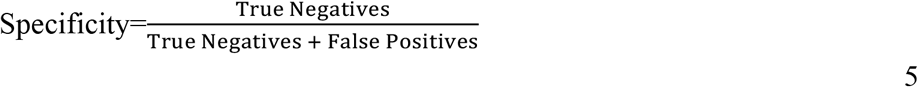

**Figure S1.**
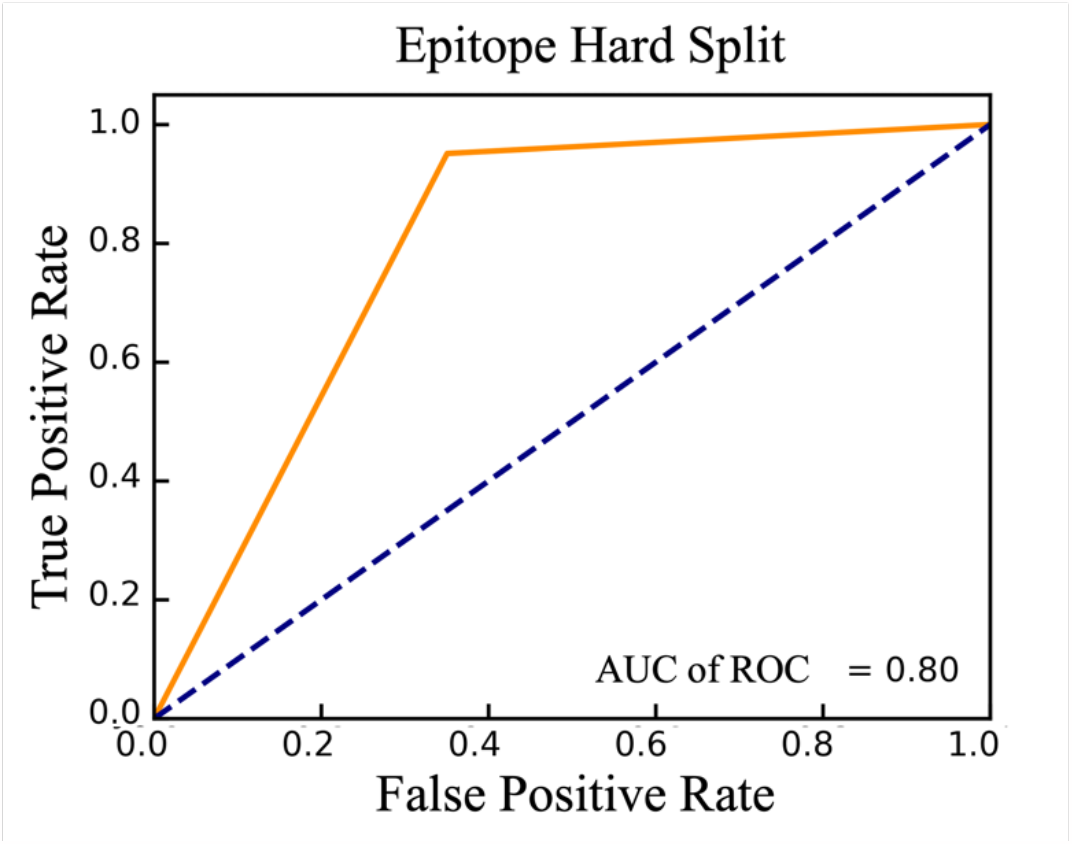
AUC of ROC on the independent test set for the SVM TCR-HE model trained with the hard split for the training and independent test set. This SVM model was trained on the set of all features, without removing correlated features.

In Table S2 we present the performance metrics of hard splits, strict splits and random splits obtained when varying the balance of positive and negative data as described in the main text.

**Table S2:** Performance metrics when varying the balance of positive and negative data.

**Epitope Hard Split:**
| AUC of ROC | TP | TN | FP | FN | Accuracy | Precision | Recall | Specificity | F1 |
| --- | --- | --- | --- | --- | --- | --- | --- | --- | --- |
| 0.853 | 12487 | 19446 | 3087 | 2300 | 0.855 | 0.801 | 0.844 | 0.863 | 0.82 |
| 0.72 | 1361 | 1624 | 1785 | 46 | 0.619 | 0.432 | 0.967 | 0.476 | 0.59 |
| 0.79 | 4470 | 164 | 91 | 294 | 0.923 | 0.98 | 0.938 | 0.64 | 0.958 |

**TCR Hard Split**
| AUC of ROC | TP | TN | FP | FN | Accuracy | Precision | Recall | Specificity | F1 |
| --- | --- | --- | --- | --- | --- | --- | --- | --- | --- |
| 0.916 | 18391 | 28487 | 765 | 3023 | 0.925 | 0.96 | 0.858 | 0.973 | 0.966 |
| 0.918 | 18462 | 28819 | 767 | 2955 | 0.927 | 0.96 | 0.862 | 0.974 | 0.908 |
| 0.918 | 18760 | 28780 | 766 | 2994 | 0.926 | 0.96 | 0.862 | 0.974 | 0.908 |

Strict Split:
| AUC of ROC | TP | TN | FP | FN | Accuracy | Precision | Recall | Specificity | F1 |
| --- | --- | --- | --- | --- | --- | --- | --- | --- | --- |
| 0.827 | 13714 | 16406 | 6127 | 1073 | 0.807 | 0.691 | 0.927 | 0.728 | 0.792 |
| 0.712 | 1355 | 1574 | 1835 | 52 | 0.608 | 0.424 | 0.963 | 0.461 | 0.589 |
| 0.777 | 4423 | 160 | 95 | 341 | 0.913 | 0.978 | 0.928 | 0.627 | 0.953 |

Random Split:
| AUC of ROC | TP | TN | FP | FN | Accuracy | Precision | Recall | Specificity | F1 |
| --- | --- | --- | --- | --- | --- | --- | --- | --- | --- |
| 0.918 | 18428 | 28762 | 700 | 2999 | 0.93 | 0.96 | 0.86 | 0.98 | 0.91 |
| 0.917 | 18358 | 28827 | 672 | 3032 | 0.93 | 0.96 | 0.86 | 0.98 | 0.91 |
| 0.917 | 18246 | 28921 | 659 | 3063 | 0.93 | 0.97 | 0.86 | 0.98 | 0.91 |

**Figure S3.**
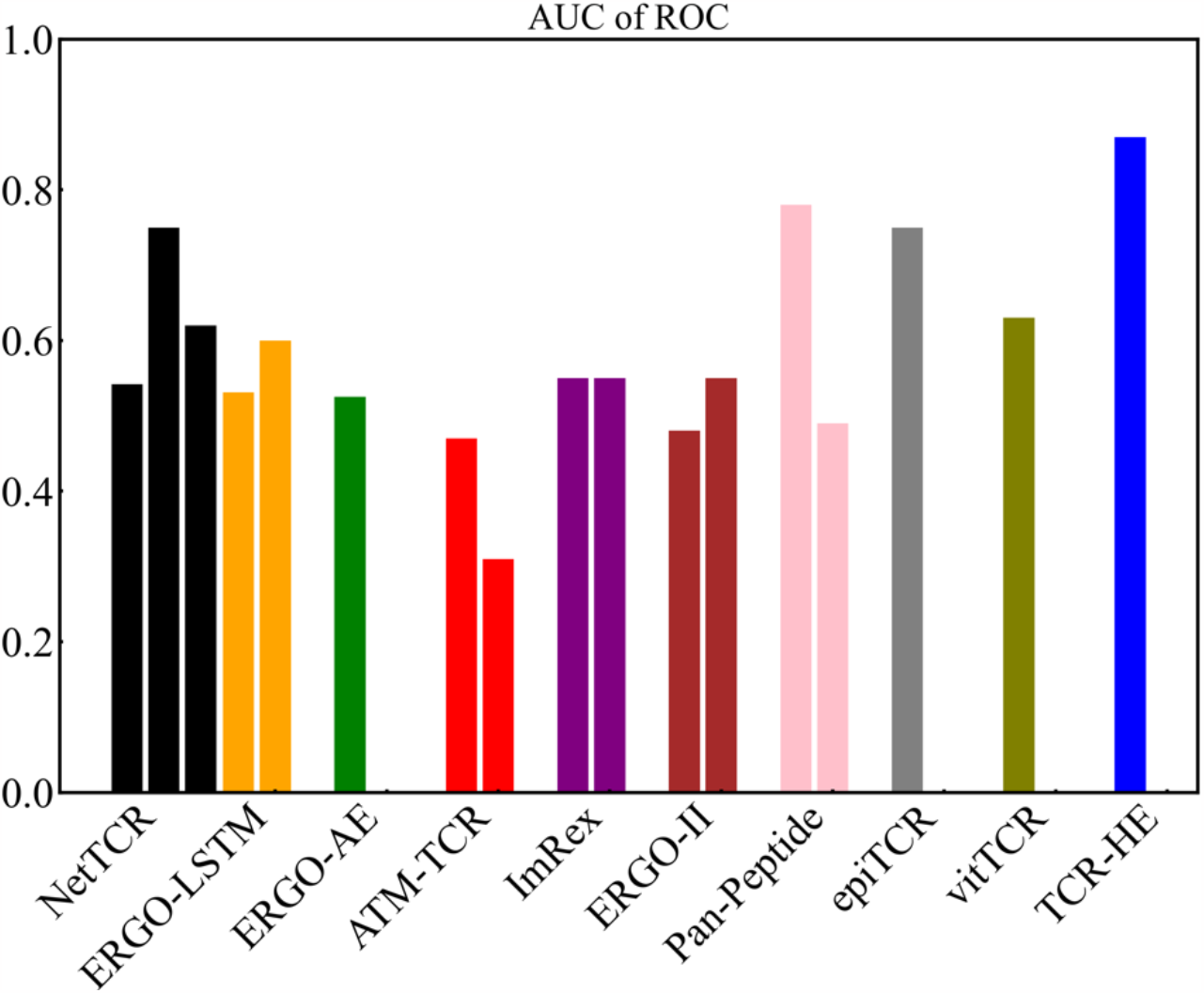
Epitope Hard Split AUC of ROC data with TCR-HE and various previously-reported models. Each bar represents the given model with the same training set but tested on a specific hard split data set. Data taken from (Cai et al., 2022; Gao, Gao, Fan, et al., 2023b; Grazioli et al., 2022; Jiang et al., 2023 biorXiv.; Pham et al., 2023)

## Notes

### Competing Interest Statement

The authors have declared no competing interest.

### Summary of Updates

Results of additional hard splits are added

